# Alpha and theta oscillations differentiate escalating risk levels during reward anticipation in sequential decision making

**DOI:** 10.1101/2025.07.08.663699

**Authors:** Eszter Tóth-Fáber, Andrea Kóbor

## Abstract

Reward anticipation potentially guides sequential decision making, yet its underlying neural dynamics remain unclear. In this study, we investigated how within-trial escalating risk and contextual uncertainty modulate oscillatory brain activity during reward anticipation. EEG was recorded while participants (N = 44) performed a modified version of the Balloon Analogue Risk Task in which balloon burst probabilities changed across task phases, introducing contextual uncertainty. Spectral power was analyzed time-locked to three within-trial risk levels: early no-risk pumps, final successful pumps (preceding cash out), and unsuccessful pumps (preceding balloon burst). Time–frequency decomposition using Morlet wavelets revealed a parieto-occipital alpha power increase following early no-risk pumps, consistent with reduced deliberative engagement when anticipating certain rewards. In contrast, centroparietal alpha suppression emerged following final successful pumps, suggesting increased attentional engagement and reward expectancy at high within-trial risk levels. Frontocentral theta power also decreased following final successful pumps, with the strongest reduction observed when the burst probability function was shallower. Rather than reflecting simple reward encoding, this pattern may index reduced monitoring demands following commitment to the selected action. Overall, both alpha and theta dynamics tracked within-trial escalating risk, whereas theta activity was additionally modulated by contextual factors across task phases. These findings provide novel insights into the oscillatory mechanisms supporting reward anticipation in sequential decision environments.

## Introduction

Reward processing is a complex neurocognitive phenomenon that unfolds across two temporally distinct stages: reward anticipation and reward outcome (e.g., Alí Diez, Fàbrega-Camps, Parra-Tíjaro, & Marco-Pallarés, 2024; Berridge, Robinson, & Aldridge, 2009; Glazer, Kelley, Pornpattananangkul, Mittal, & Nusslock, 2018; Meyer, Marco-Pallarés, Boulinguez, & Sescousse, 2021). While extensive research has examined the neural mechanisms underlying reward outcome, the spectral EEG dynamics associated with reward anticipation remain relatively underexplored (for review, see Meyer et al., 2021). Understanding these dynamics is crucial, as reward anticipation plays a key role in guiding decision making, particularly in uncertain environments where learning from and predicting probabilistic rewards are essential (Alí Diez et al., 2024; Berridge & Robinson, 2003).

Most of our daily decisions take place under uncertain conditions, requiring us to gradually learn the probabilities of different outcomes (De Groot & Thurik, 2018; Hertwig & Erev, 2009; Knight, 1921). For example, when playing an arcade claw machine game with a limited number of coins, we make repeated decisions on whether to continue playing based on previous successes or failures. Each time we insert a coin and control the claw, we anticipate whether we will successfully grab a prize or not. If the claw has weak grip strength multiple times in a row, we might expect failure and hesitate to keep playing. Conversely, a series of successful or nearly successful grabs might encourage further attempts. This sequential decision-making process mirrors how individuals adjust expectations and risk-taking behavior in uncertain situations. Yet, it remains unclear whether changes in contextual uncertainty and escalating risk within a decision sequence shape anticipatory neural activity at the time–frequency level. To bridge this research gap, we investigate how these factors jointly influence EEG spectral dynamics during reward anticipation following a decision in a task simulating real-life risk taking.

To this end, we use a modified version of the Balloon Analogue Risk Task (BART), a well-established paradigm for studying naturalistic risk-taking behavior (De Groot, 2020; Helfinstein et al., 2014; Lejuez et al., 2002; Schonberg, Fox, & Poldrack, 2011). In this task, participants make sequential decisions to inflate a virtual balloon, thereby increasing the potential rewards while also increasing the a priori unknown probability of the balloon bursting, which would result in the loss of accumulated earnings. Classical decision theory distinguishes risk (known probabilities) from ambiguity (unknown probabilities) (De Groot & Thurik, 2018; Hertwig & Erev, 2009; Jessup, Busemeyer, Dimperio, Homer, & Phillips, 2022). Because burst probabilities are never disclosed to participants, the present BART variant constitutes an experience-based ambiguity task (e.g., De Groot, 2020; De Groot & Thurik, 2018; Fecteau et al., 2007).

Previous event-related brain potential (ERP) studies have explored reward processing within the BART, focusing primarily on the outcome stage (e.g., Gu, Zhang, Luo, Wang, & Broster, 2018; Kardos et al., 2016; Kiat, Straley, & Cheadle, 2016; Kóbor et al., 2015; Petropoulos Petalas, Bos, Hendriks Vettehen, & van Schie, 2020; Xu, 2021). Some of these studies have demonstrated progressive expectation formation about the reward structure of the task (Kardos et al., 2016; Kiat et al., 2016; Petropoulos Petalas et al., 2020). However, the investigation of reward expectation and anticipation was conducted indirectly through the analysis of outcome-related processing at the various steps of the balloon inflation process using stimulus-locked ERPs. Consequently, the EEG dynamics related to each pump during balloon inflation, while awaiting and anticipating the outcome (inflated balloon or balloon burst), remained to be elucidated. The latter approach, which is also employed in this study, has the potential to directly reveal whether reward expectation and anticipation undergo changes over the course of the task.

Reward expectation and anticipation can change dynamically as the reward structure of the decision environment shifts. Prior BART studies have demonstrated that participants learn burst probability functions over time and adapt their behavior accordingly (Bonini, Grecucci, Nicolè, & Savadori, 2018; Kóbor et al., 2021; Kóbor et al., 2023; Koscielniak, Rydzewska, & Sedek, 2016; Smith, Ebert, & Broman-Fulks, 2016; Young & McCoy, 2019). Building on this evidence, the present study therefore employs a three-phase manipulation – baseline, lucky, and unlucky – to systematically alter the burst probability structure of the task. The lucky phase is characterized by a shallower increase in burst probability, whereas the unlucky phase involves a steeper hazard trajectory relative to baseline.

Importantly, computational learning frameworks distinguish between expected uncertainty – irreducible stochasticity within a stable generative model – and unexpected uncertainty, which arises from unsignaled changes in that model (i.e., environmental volatility) (Soltani & Izquierdo, 2019; Yu & Dayan, 2005). In the present design, stochastic burst outcomes within each phase reflect expected uncertainty under stable burst probability functions. In contrast, the unsignaled transitions between phases introduce contextual uncertainty consistent with unexpected uncertainty, as they require participants to infer shifts in the underlying burst probability structure through experience. We therefore distinguish between *within-trial escalating risk* – reflecting conditional hazard increases across inflation steps within a balloon – and *contextual uncertainty* across phases.

We have previously shown that the three-phase manipulation (baseline, lucky, unlucky) influences how individuals learn from rewards, as revealed by computational modeling and behavioral indices (Bán, Tóth-Fáber, Lages, & Kóbor, 2024). Specifically, heightened sensitivity to rewards relative to losses was associated with heightened risk-taking behavior across all phases. Moreover, this heightened sensitivity was inversely correlated with task performance in the unlucky phase, which was characterized by a steeper burst probability function and elevated conditional hazard. In the present study, we extend these findings by examining the spectral EEG components associated with reward anticipation that support processing across both within-trial escalating risk levels and contextual uncertainty introduced by phase-wise changes in burst probability.

EEG time–frequency measures provide several advantages over traditional ERP analyses. Unlike ERPs, which primarily reflect phase-locked activity, time–frequency analyses can also capture non-phase-locked oscillatory dynamics, providing a richer representation of neural activity (Alí Diez et al., 2024; Bastiaansen, Mazaheri, & Jensen, 2012; Cohen, 2014; Glazer et al., 2018; Li, Baker, Warren, & Li, 2016; Makeig et al., 2002; Tallon-Baudry & Bertrand, 1999). These measures can allow us to investigate the fine-grained temporal dynamics of anticipatory neural activity. So far, with time–frequency analysis, research on sequential risk-taking using variants of the BART has targeted only the processing of reward outcomes. In the outcome stage, midfrontal theta activity has been shown to vary depending on outcome type (Edgar et al., 2024), personality traits related to risk taking (Edgar et al., 2024; Qianlan, Shou, Tianya, Wei, & Liu, 2025), and gain/loss framing (Tao et al., 2023). To our knowledge, no studies have investigated time–frequency dynamics during reward anticipation in this task.

In the context of reward anticipation, characteristics of EEG time–frequency components have been predominantly examined within paradigms that diverge from the BART, including monetary incentive delay and time-estimation tasks with cue-target-outcome structures. In this earlier research, parieto-occipital alpha suppression has been found to reflect increased attention allocation and disinhibition of sensory cortices in preparation for outcome processing (Bastiaansen, Böcker, & Brunia, 2002; Bastiaansen & Brunia, 2001; Gómez, Vaquero, López-Mendoza, González-Rosa, & Vázquez-Marrufo, 2004; Pornpattananangkul & Nusslock, 2016; van den Berg, Krebs, Lorist, & Woldorff, 2014; Zhang et al., 2023). This suppression is typically greater before rewarding than nonrewarding outcomes as well as before high than low magnitude outcomes, suggesting its role in facilitating adaptive behavior. Additionally, anterior beta activity has also been associated with reward anticipation, often increasing before higher potential rewards, indicative of incentive-motivational processes (Apitz & Bunzeck, 2014; Bunzeck, Guitart-Masip, Dolan, & Düzel, 2011; Reinhart & Woodman, 2014; Xiao et al., 2024). However, findings regarding anticipatory frontocentral theta activity have remained mixed. Some studies report increased theta power prior to (higher) potential rewards (Alí Diez et al., 2024; Fryer et al., 2021; Reinhart & Woodman, 2014; Zhang et al., 2023), whereas others observe a decrease (Bunzeck et al., 2011; Doñamayor, Schoenfeld, & Münte, 2012) or no reliable modulation as reward probability increases (Lopez-Gamundi, Mas-Herrero, & Marco-Pallares, 2024). The general reduction of EEG power during expectancy periods has also been noted in a nonrewarded cue-target paradigm, further emphasizing the complexity of anticipatory neural dynamics (Gómez et al., 2004).

Building on this background, we examine how different levels of within-trial escalating risk and contextual uncertainty influence EEG spectral activity during reward anticipation in the BART. We leverage previously established experimental contrasts (Kardos et al., 2016) that capture reward expectation evolving throughout the balloon inflation process. These contrasts define the key within-trial risk levels that we analyze in this study (see Figure 1). The first level is the early no-risk pump that serves as a zero-hazard reference condition within the escalating conditional risk structure: a sure reward (inflated balloon) can be anticipated after this pump (second pump; burst probability = 0). The second level is the final pump prior to cashing out. Due to the success of this final pump, a reward occurs eventually. The third level is the pump leading to a balloon burst. The second and third levels are located at the end of the balloon trial, and subsequent to both of these pumps, either a balloon burst or a reward may be anticipated. Critically, the outcome remains unpredictable until the moment it occurs. According to the findings of ERP and fMRI studies conducted previously, there is an indication that brain responses might vary prior to cashing out as opposed to pumping towards a balloon burst (i.e., reward positivity and P3 amplitudes, cognitive control networks, see Helfinstein et al., 2014; Kardos et al., 2016). Consequently, the question remains open regarding whether and, if so, how anticipation differs between the final successful and unsuccessful pumps. In addition to these risk levels, we examine the effect of contextual uncertainty introduced by the three-phase task structure (baseline, lucky, unlucky). Note that participants were not informed about burst probabilities, the changes in these probabilities across task phases, and the zero probability of balloon burst after the initial two pumps.

**Figure 1.**
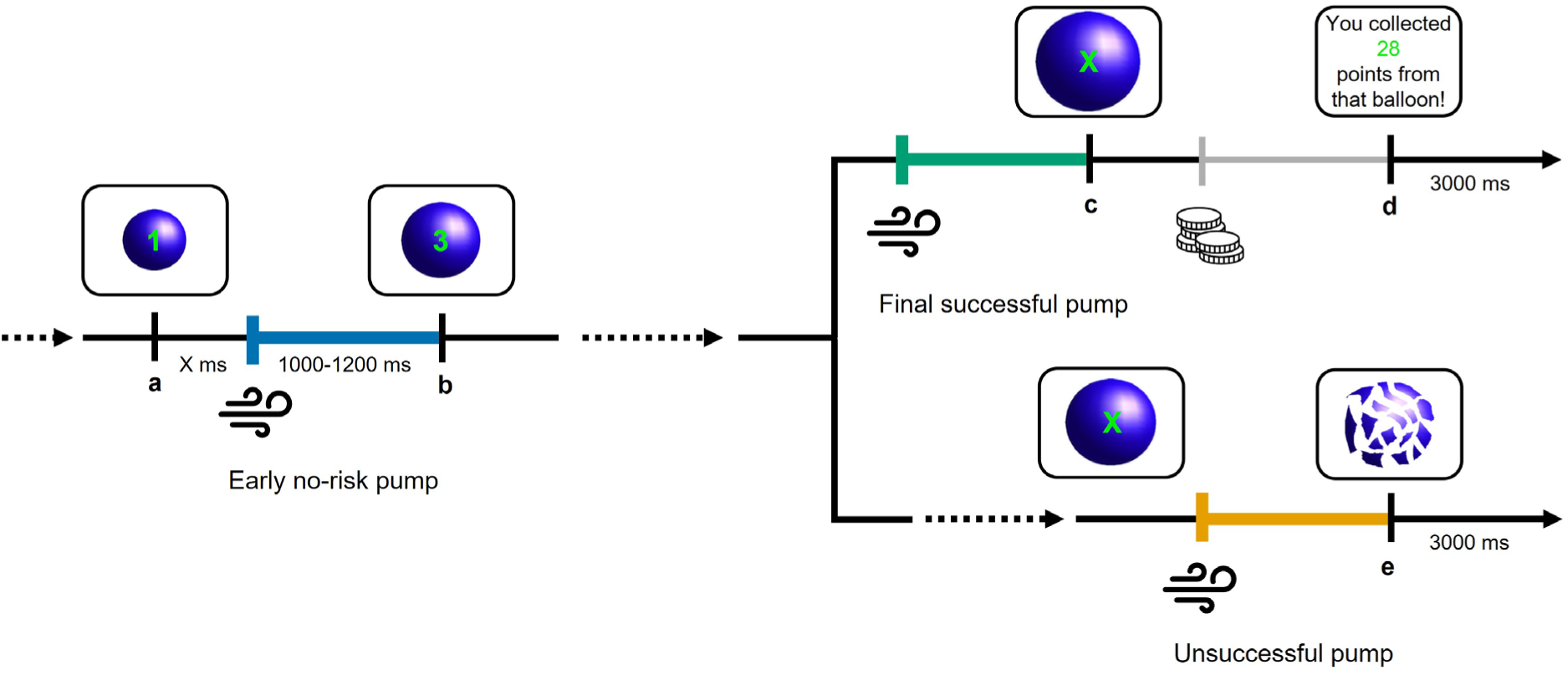
Schematic illustration of the trial structure and the key within-trial risk levels in the BART. Each balloon trial started with the presentation of a zero-value balloon (not indicated in this figure). Lowercase capital letters (a, b, c, d, e) denote the onsets of balloon presentation after the first (a), second (b), and further (c) pumps, and the onsets of feedback presentation at the end of the balloon trial (d = cash-out screen for successful balloon trials, e = balloon burst for unsuccessful balloon trials). The blow icons represent the onset of the pump response, relative to which the EEG segments were synchronized in time. The coins icon represents the onset of the cash-out response, by which participants could collect the accumulated score from the current balloon and end the balloon trial. The score accumulated in the current balloon trial was presented in the middle of the balloon during the task. Participants had no time limit (“X ms”) to respond to the current balloon, i.e., to pump the balloon further or to cash out. After each successful pump, one point more was added to the accumulated score (1, 3, 6, 10, 15, etc., presented as the accumulated score). Each response was followed by the inflated balloon stimulus (reward) or the end-of-trial feedback stimulus (cash-out screen or burst stimulus) after a random delay of 1000-1200 ms. This delay period was considered as the reward anticipation stage, where EEG spectral dynamics were analyzed by time–frequency analysis. Note that the current balloon was presented on the screen throughout the response and delay periods (i.e., until the onset of the next balloon or that of the feedback). The blue-, green-, and yellow-colored, thick sections of the timeline denote the reward anticipation stages. The blue section is the anticipatory stage after the early no-risk (second) pump, the green section is the anticipatory stage after the final successful pump before cashing out, and the yellow section is the anticipatory stage after the unsuccessful pump leading to a balloon burst. The occurrence of the final successful pump, as well as the unsuccessful pump, is possible at any subsequent time point (denoted by dashed arrow lines) after the early no-risk pump. Therefore, the exact value of the balloon is not given for these risk levels (indicated by green “X”). The anticipatory stage after the cash-out response (grey-colored, thin section) was not analyzed. For more details on the task structure and the expectedness of the reward, see the main text. This illustration has been adapted from the figure published by Kardos et al. (2016).

Regarding the within-trial risk levels, we hypothesize that alpha suppression will become more pronounced as conditional hazard increases across successive inflation steps, with the largest suppression following both successful and unsuccessful pumps at the final inflation step. At the early no-risk inflation step, alpha power may instead increase, reflecting neural inhibition necessary for continuing the inflation process without deliberation (Klimesch, 2012). Beta power is expected to be modulated by outcome probability within the escalating hazard structure, increasing when reward probability is higher at the early no-risk step. Furthermore, given reports that anticipatory theta can decrease when reward probability is higher (Bunzeck et al., 2011; Doñamayor et al., 2012), we tentatively expect theta suppression at the early no-risk inflation step and relatively increased theta at the final inflation steps, where outcome uncertainty increases. Beyond reward probability, the magnitude of the potential reward could also induce theta enhancement at the final inflation steps relative to the early no-risk step (Zhang et al., 2023). Alpha, beta, and theta power may differentiate between successful and unsuccessful balloon inflations; however, this remains an exploratory question (Helfinstein et al., 2014; Kardos et al., 2016).

Regarding contextual uncertainty, we predict that alpha suppression will be strongest in the unlucky phase, which is characterized by a steeper burst probability function and elevated conditional hazard. Beta power is expected to be higher in phases with more probable rewards (i.e., the lucky and baseline phases). To adapt to altered reward-loss contingencies in the lucky and unlucky phases, enhanced top-down control may be required. Accordingly, theta power may increase in these phases relative to the baseline phase (Zhang et al., 2023). In the unlucky phase, increased theta may reflect heightened performance monitoring demands under elevated loss probability, whereas in the lucky phase, it may index greater effort investment and sustained reward pursuit (Lopez-Gamundi et al., 2024). Finally, we hypothesize an interaction between within-trial escalating risk and contextual uncertainty, such that the predicted time–frequency modulations across risk levels will be most pronounced under elevated burst probability (i.e., in the unlucky phase). Through these analyses, we aim to provide novel insights into the spectral dynamics of reward anticipation under conditions of escalating within-trial risk and phase-wise contextual uncertainty. This could advance our understanding of neurocognitive mechanisms supporting sequential decision making.

## Materials and methods

### Participants

Fifty young adults took part in the experiment. They were recruited from university courses and were randomly assigned to one of the two order conditions (about the order, see Task section). All participants had normal or corrected-to-normal vision, reported no history of psychiatric or neurological disorders, and were not taking any psychoactive medication at the time of participation. Handedness was assessed using the revised version of the Edinburgh Handedness Inventory (Dragovic, 2004a, 2004b; Oldfield, 1971). The mean laterality quotient was 75.71 (*SD* = 40.68; −100 means complete left-handedness, 100 means complete right-handedness). During the EEG analyses, six participants were excluded due to excessive artifacts: For them, the mean ratio of the removed EEG segments was greater than 25% (for details, see EEG recording and analyses section). Hence, the final sample consisted of 44 participants (*M_age_* = 21.23 years, *SD_age_* = 2.52 years, 8 males, 36 females). A detailed justification of the sample size is provided in the Supplementary Materials (see section Justification of the sample size).

Prior to enrollment, all participants provided written informed consent. The study received ethical approval from the United Ethical Review Committee for Research in Psychology (EPKEB) in Hungary and was conducted in accordance with the principles of the Declaration of Helsinki. In exchange for their participation, students received course credit and a supermarket voucher. Although the participants were informed that the value of the voucher would depend on their task performance (ranging from 1000 to 2000 HUF, approximately €2.5– €5), all received a voucher worth 2000 HUF upon completion of the experiment.

### Task

A modified version of the BART (Bán et al., 2024; Fein & Chang, 2008; Kóbor et al., 2015; Kóbor et al., 2023) was employed in the present study. During the task, participants were instructed to collect as many points as possible by inflating virtual balloons on the screen, one at a time. They were required to repeatedly decide whether to continue inflating a balloon or to stop and collect their accumulated score.

Each successful pump increased both the balloon’s size and the potential reward, while simultaneously raising the probability of the balloon bursting. Participants were instructed to use two designated response keys to indicate their decision either to pump the balloon further or to terminate the trial and collect the accumulated score. Each inflation could result in one of two outcomes: either the balloon successfully inflated and the score increased, or the balloon burst, resulting in the loss of the trial’s accumulated score. If participants decided to stop inflating the balloon, the score collected during that trial were transferred to a virtual permanent bank. Bursting a balloon did not affect the score stored in the permanent bank.

Throughout the task, the current trial’s accumulated score was displayed within the balloon, while the permanent bank score, the score collected in the previous trial, and the response options for pumping or cashing out were continuously visible on the screen. A balloon burst was visually indicated by a fragmented balloon stimulus, whereas cash-out trials were followed by a feedback screen displaying the score collected on that balloon trial. The duration in which a response (i.e., pump or cash out) could be initiated was unrestricted. Each response was followed by the inflated balloon stimulus (reward) or the end-of-trial feedback stimulus (cash-out screen or burst stimulus) after a random delay of 1000-1200 ms. Feedback for both cash-out and burst trials was presented for 3000 ms (Figure 1).

The task was structurally divided into three phases, each corresponding to different balloon burst probabilities. The baseline phase featured an intermediate probability of bursting. This was followed by either an increase (unlucky phase) or a decrease (lucky phase) in burst probability. All participants completed the baseline phase first. Subsequently, half were assigned to complete the lucky phase followed by the unlucky phase (Lucky-first order), while the remaining participants completed the unlucky phase first and concluded with the lucky phase (Unlucky-first order) (see also Bán et al., 2024).

In all phases, balloon bursts were disabled for the first and second pumps. In the baseline phase, the burst probability began at 1/18 on the third pump and increased gradually (1/17 on the fourth pump, 1/16 on the fifth, etc.), reaching a certainty of bursting (probability = 1) on the 20th pump. In the lucky phase, the burst probability followed a less steep trajectory (1/28 on the third pump, 1/27 on the fourth pump, 1/26 on the fifth, etc.) but burst probability was set to 1 on the 20th pump. This design was chosen to maintain a manageable average trial duration. In contrast, the unlucky phase was characterized by a steeper increase in burst probability (1/8 on the third pump, 1/7 on the fourth pump, 1/6 for the fifth, etc.), with the maximum burst probability reached on the 10th pump. In other words, burst probability for a given pump was defined by the following truncated power function: *p*_burst_ = (tolerance + 2 − pump_*n*_)^−1^, where pump*_n_* is the n^th^ pump (*n* ≥ 3) on a given balloon and tolerance is equal to 19, 29, and 9 in the baseline, lucky, and unlucky phases, respectively, with the restriction that burst probability was a priori fixed as 1 at the 20th pump in the lucky phase. For more details, see Figure 2 in Bán et al. (2024) and Tables S2-S4 in the Supplementary Methods of Kóbor et al. (2023). Participants were naïve regarding the burst probabilities in the experiment, including the zero probability of balloon burst in the first two inflation opportunities. Participants were also unaware that burst probabilities would change during the experiment, and the beginning of a new phase was unsignaled.

**Figure 2.**
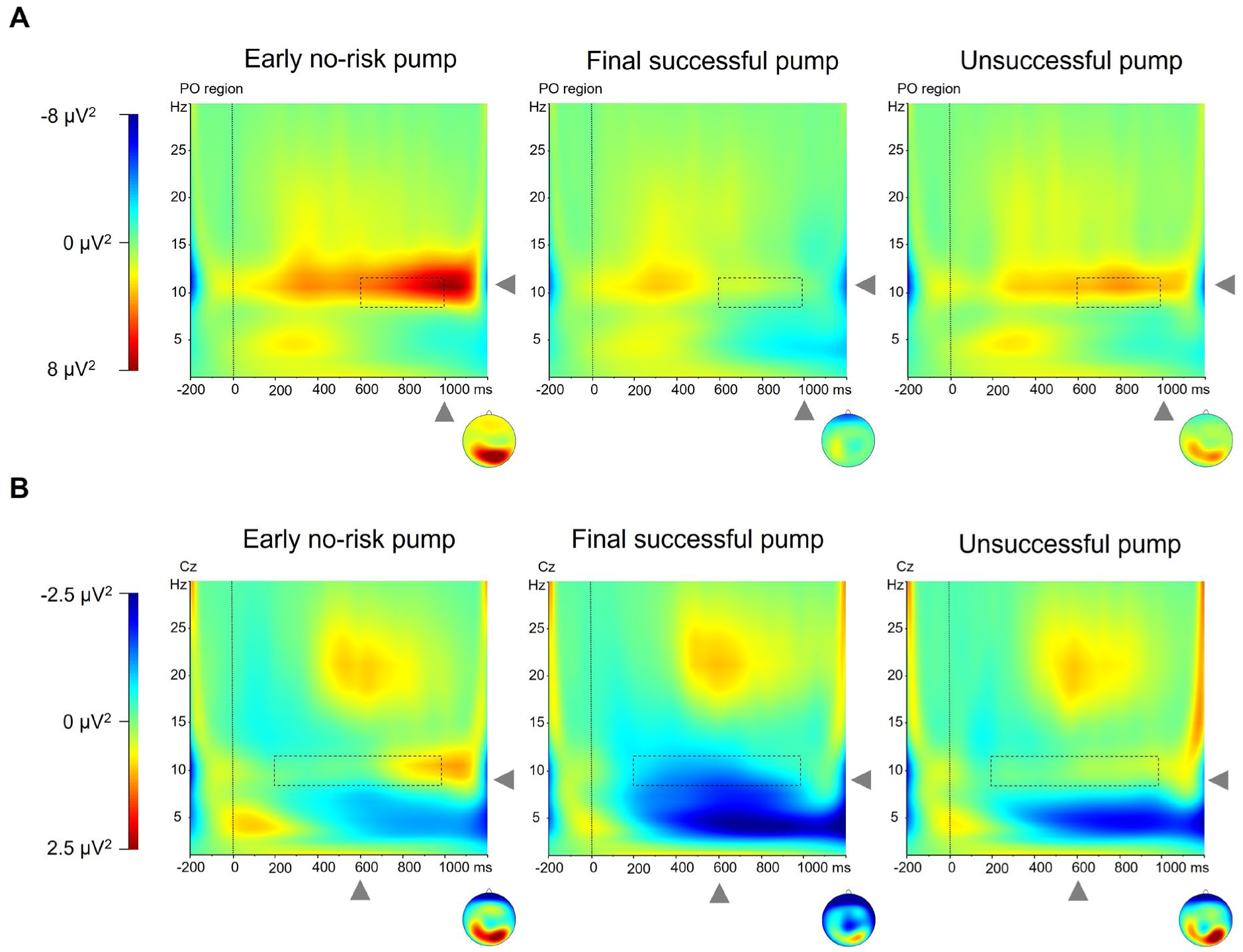
Time–frequency representations of total alpha power across outcomes. **(A)** Grand-average time–frequency plots of alpha power over the parieto-occipital (PO) region (O1, O2, Oz, POz) across the three outcome types. The time–frequency window used for analysis (600–1000 ms post-response, 8.26 Hz to 11.74 Hz) is shown with dashed rectangles. A prominent increase in alpha power was observed after the early no-risk pumps. **(B)** Grand-average time–frequency plots of alpha power at Cz across the three outcome types. The time–frequency window used for analysis (200–1000 ms post-response, 8.26 Hz to 11.74 Hz) is shown with dashed rectangles. A prominent decrease in alpha power was observed after the final successful pumps. Topographical maps to the right of each plot illustrate the scalp distribution of alpha power at the corresponding time–frequency point (indicated by the grey arrowheads). The response (pump) onset was at 0 ms.

Each phase consisted of 90 balloon trials, out of which participants could freely decide whether they inflate the balloon or collect their accumulated score in 80 trials. In 10 trials of each phase, participants were instructed to inflate the balloon to a predetermined number of pumps or until it burst. These forced-choice trials were incorporated to provide guidance regarding the optimal number of pumps in each phase. To minimize inter-individual variability, the position of these trials was fixed and identical across all participants.

The task was implemented using Presentation software (Version 21.1; Neurobehavioral Systems, Inc., Berkeley, CA), and participants’ responses were recorded via a Cedrus RB-540 response device (Cedrus Corporation, San Pedro, CA).

### Procedure

Participants completed two experimental sessions. The first session involved a series of well-established neuropsychological assessments targeting executive functions, aimed at confirming that all participants performed within the neurotypical range. This session lasted approximately one hour. The second session consisted of the BART, which included both a practice and a test session. During practice, participants completed six forced-choice trials to ensure a standardized initial experience with the task. The test session consisted of 270 balloon trials (3*90) in total. To reduce fatigue, brief breaks were incorporated every 20–25 trials, with a longer break scheduled at the midpoint of the task. Continuous EEG data were recorded throughout the session. Upon completion of the BART, participants took part in a brief verbal interview, during which they were asked about their strategies and whether they had noticed any changes in burst probabilities or differences across the task phases. The second session lasted approximately 2 to 2.5 hours, including the application and removal of the electrode cap.

Verbal interviews were recorded using a portable audio recording device (OLYMPUS WS-852). The recordings were later evaluated by an independent rater who was blind to both the research hypotheses and the experimental conditions. A bottom-up, data-driven approach was employed for the initial categorization: Participants were grouped based on the similarity of the patterns they reported noticing during the task. These groups were then categorized according to whether participants had detected the full extent of the phase-wise experimental manipulation, only partial aspects of it, or no regularities at all. Following this initial grouping, the rater was debriefed about the nature of the experimental manipulation to inform final categorization. Among the 44 participants in the final sample, 19 (43.2%) reported noticing changes consistent with the structure of the task phases, 16 (36.4%) noticed some phase-related changes without fully grasping the manipulation, and 9 (20.5%) did not report noticing any changes. Beyond the phase-wise changes, participants were unable to report the burst probability functions governing balloon explosions.

### EEG recording and analyses

The continuous EEG activity was recorded in an electrically shielded, acoustically attenuated, and dimly lit room using the actiCAP slim/snap active electrode system with BrainAmp Standard amplifier and BrainVision Recorder software (Version 1.2; BrainProducts GmbH, Gilching, Germany). EEG signals were acquired from 64 Ag/AgCl electrodes with integrated active circuits, mounted in an elastic cap and positioned according to the 10% equidistant system. The FCz electrode served as the reference, while Fpz was used as the ground. The sampling rate was 1000 Hz. Electrode impedance was maintained below 10 kΩ throughout recording.

Offline EEG processing was conducted using BrainVision Analyzer software (Version 2.2.2; BrainProducts GmbH). First, the continuous EEG data were visually inspected for major artifacts: When necessary, noisy electrodes were corrected via spherical spline interpolation, with a maximum of two interpolated electrodes per participant (*M* = 0.23, *SD* = 0.47). Second, ocular and cardiac artifacts were removed using independent component analysis (ICA; Delorme, Sejnowski, & Makeig, 2007). Between two and four components per participant were rejected (*M* = 2.72, *SD* = 0.54), after which the channel-based EEG data were recomposed. Third, the data were band-pass filtered between 0.03–30 Hz (48 dB/oct) and notch filtered at 50 Hz to remove additional electrical noise. Fourth, the data were re-referenced to the average of all electrodes, including FCz, which had served as the online reference. The FCz was then reused in the subsequent analyses. Finally, the continuous, preprocessed EEG signal was segmented according to the criteria outlined below.

Based on the within-trial escalating risk structure and the phase manipulation, three outcome-defined event types were extracted for EEG segmentation (see Figure 1). Although defined by eventual outcome, these event types correspond to distinct positions within the conditional hazard structure and therefore index different within-trial risk levels. For clarity, we refer to them as outcome types in the statistical analyses, while interpreting them conceptually as reflecting differences in within-trial risk. The *early no-risk pump* was defined as the second pump of each balloon trial, during which a balloon burst was experimentally disabled. These pumps were associated with the anticipation of a guaranteed reward, as each pump was certain to result in the inflation of the balloon and an increase in the temporary score. The *final successful pump* referred to the last pump initiated before the participant chose to cash out. The *unsuccessful pump* was defined as the pump immediately preceding a balloon burst, resulting in the loss of the accumulated score. At these points in the balloon trial, the outcome remained uncertain—either a balloon burst or a reward could occur. Therefore, it can be posited that these pumps were associated with the anticipation of both outcomes.

Each response in the task was followed by a random delay of 1000-1200 ms before presenting its result (i.e., reward/inflated balloon, cash-out screen, or balloon burst, see Figure 1). EEG segments were time-locked to the pump responses that preceded the three outcome types of interest: early no-risk pumps (second pumps, when burst was disabled), final successful pumps (the last pumps before a cash out), and unsuccessful pumps (pumps that triggered a balloon burst). For each of these outcome types, segments were extracted from −200 ms to 1200 ms relative to response onset. These outcome types were extracted separately for each phase of the task (baseline, lucky, and unlucky), resulting in altogether nine experimental conditions. Note that, although outcome types have been identified, they should not be confused with the potential stimulus-locked analysis of actual response outcomes.

Following segmentation, to remove remaining artifacts after ICA corrections, an automatic artifact rejection algorithm implemented in the BrainVision Analyzer software rejected segments where the activity exceeded ± 100 μV at any of the electrode sites. Across the three phases, the mean numbers of the retained (artifact-free) segments were 74.40 (*SD* = 4.27) for the early no-risk pumps, 33.28 (*SD* = 8.19) for the unsuccessful pumps, and 41.05 (*SD* = 6.81) for the final successful pumps. The mean proportion of the removed segments was 6.29% (*SD* = 5.18%) for the early no-risk pumps, 5.35% (*SD* = 5.71%) for the unsuccessful pumps, and 4.43% (*SD* = 4.08%) for the final successful pumps.

Time–frequency decomposition of the retained EEG segments was conducted using a complex Morlet wavelet transform with a Morlet parameter of 5. Frequencies ranged from 1 to 30 Hz, sampled in 30 logarithmically spaced steps. Instantaneous amplitude (Gabor normalization) was used for wavelet normalization, and the resulting spectral power was extracted as real values in units of µV² (Cohen, 2014). Only artifact-free segments were included, which were baseline-corrected using the –200 to 0 ms interval preceding pump onset. For each participant, the total power was quantified as the average value of time–frequency power across single segments; then, grand averages of the total power across participants were created separately for each combination of outcome and phase (i.e., the nine experimental conditions) (Cohen, 2014; Li et al., 2016; Tallon-Baudry & Bertrand, 1999).

We examined whether oscillatory activity in the frequency ranges outlined in the Introduction (alpha, beta, and theta) varied as a function of within-trial risk levels and task phases. We considered both experimental manipulations to the same degree. Electrode locations and time windows of interest were defined based on a combination of prior literature and the overall patterns observed in the grand-average time–frequency representations (see Figures 2-3). Although time–frequency activity extended beyond 1000 ms, analyses were restricted to the 0–1000 ms post-response interval to avoid contamination from neural responses related to the subsequent stimulus presentation.

**Figure 3.**
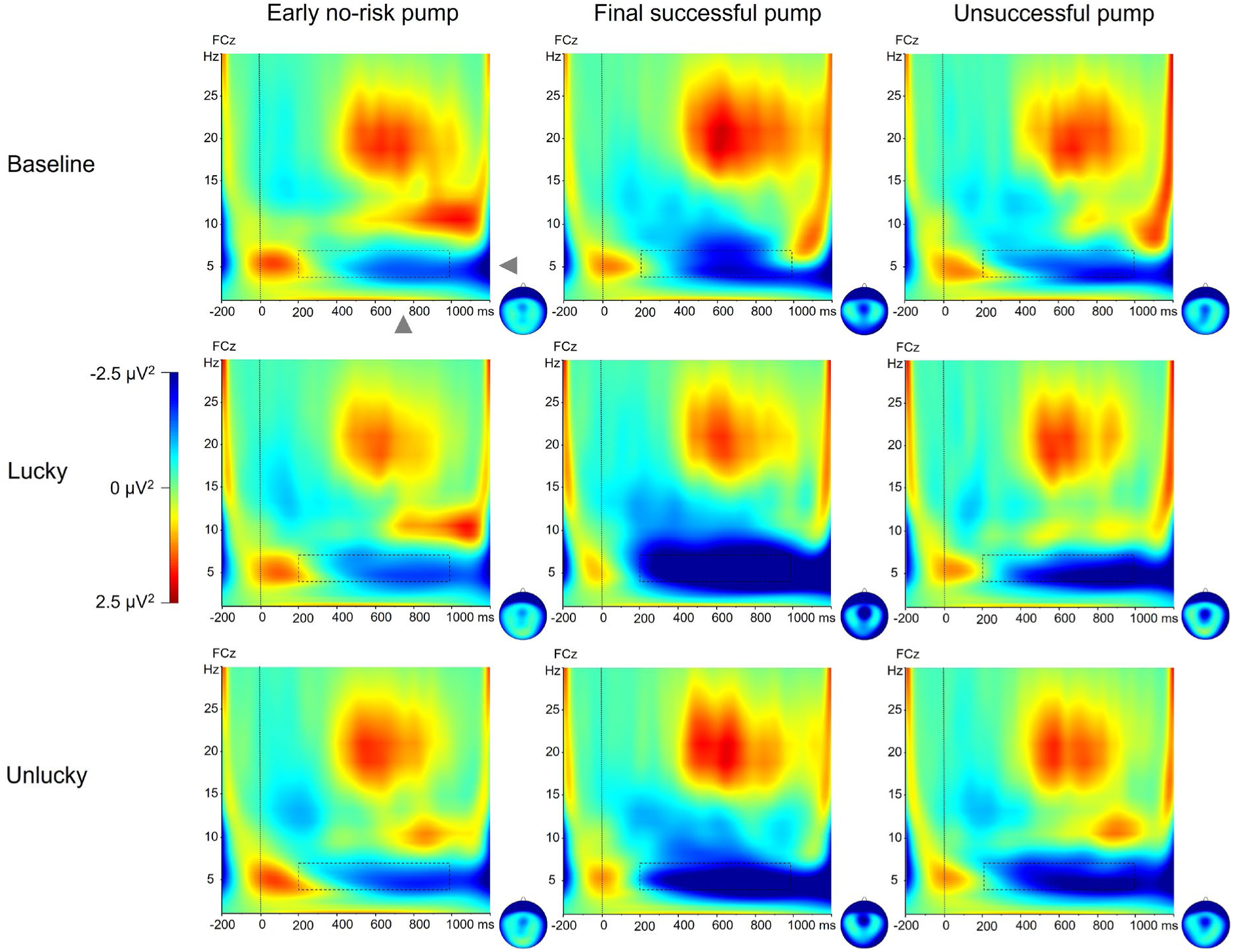
Time–frequency representations of total theta power across task phases and outcomes. Grand-average time–frequency plots of theta power at FCz are displayed for each combination of task phase (baseline, lucky, unlucky) and outcome (early no-risk pump, final successful pump, unsuccessful pump). The time–frequency window used for analysis (200–1000 ms post-response, 4.09 Hz to 7.34 Hz) is shown with dashed rectangles. Theta power decreased most markedly after the final successful pumps, especially during the lucky phase. Topographical maps to the right of each plot illustrate the scalp distribution of theta power at the corresponding time–frequency point (indicated by the grey arrowheads). The response (pump) onset was at 0 ms.

In the alpha band, two qualitatively distinct patterns were identified. First, an increase in alpha power was observed between 600 and 1000 ms over parieto-occipital sites following early no-risk pumps (Figure 2A). Mean spectral power (8.26–11.74 Hz) was therefore extracted from this time window and averaged across a posterior region of interest comprising O1, O2, Oz, and POz electrodes. Second, a decrease in alpha power emerged following final successful pumps, beginning approximately 200 ms after response onset and extending up to 1000 ms, with a focal central–centroparietal distribution (Figure 2B). To capture this effect, mean spectral power (8.26–11.74 Hz) was extracted from this time window at the Cz electrode, where the effect was most pronounced and showed the clearest selectivity across the within-trial risk levels. Neighboring posterior electrodes did not show comparable suppression, indicating that this effect was spatially distinct from the parieto-occipital alpha increase.

In the theta band, a decrease in power was observed following final successful pumps between 200 and 1000 ms over frontocentral sites (Figure 3). Accordingly, mean spectral power (4.09–7.34 Hz) was extracted from this time window at the FCz electrode. In the beta band, no consistent modulations were observed as a function of within-trial risk level or task phase in the time–frequency representations. Nevertheless, based on theoretical considerations, mean spectral power was extracted in the 16.69–23.73 Hz range between 500 and 1000 ms over a left centroparietal region of interest (C3, C5, CP3, CP5), defined based on the overall scalp distribution that was similar across the conditions.

### Statistical analysis

The statistical analysis was performed using R 4.5.0. Analyses of variance (ANOVAs) were conducted using the *aov* function from the *stats* package, and post-hoc pair-wise comparisons were carried out using estimated marginal means implemented in the *emmeans* package. Figure 4 and Figure S1 were created with the *ggplot2* package (Wickham, 2016).

**Figure 4.**
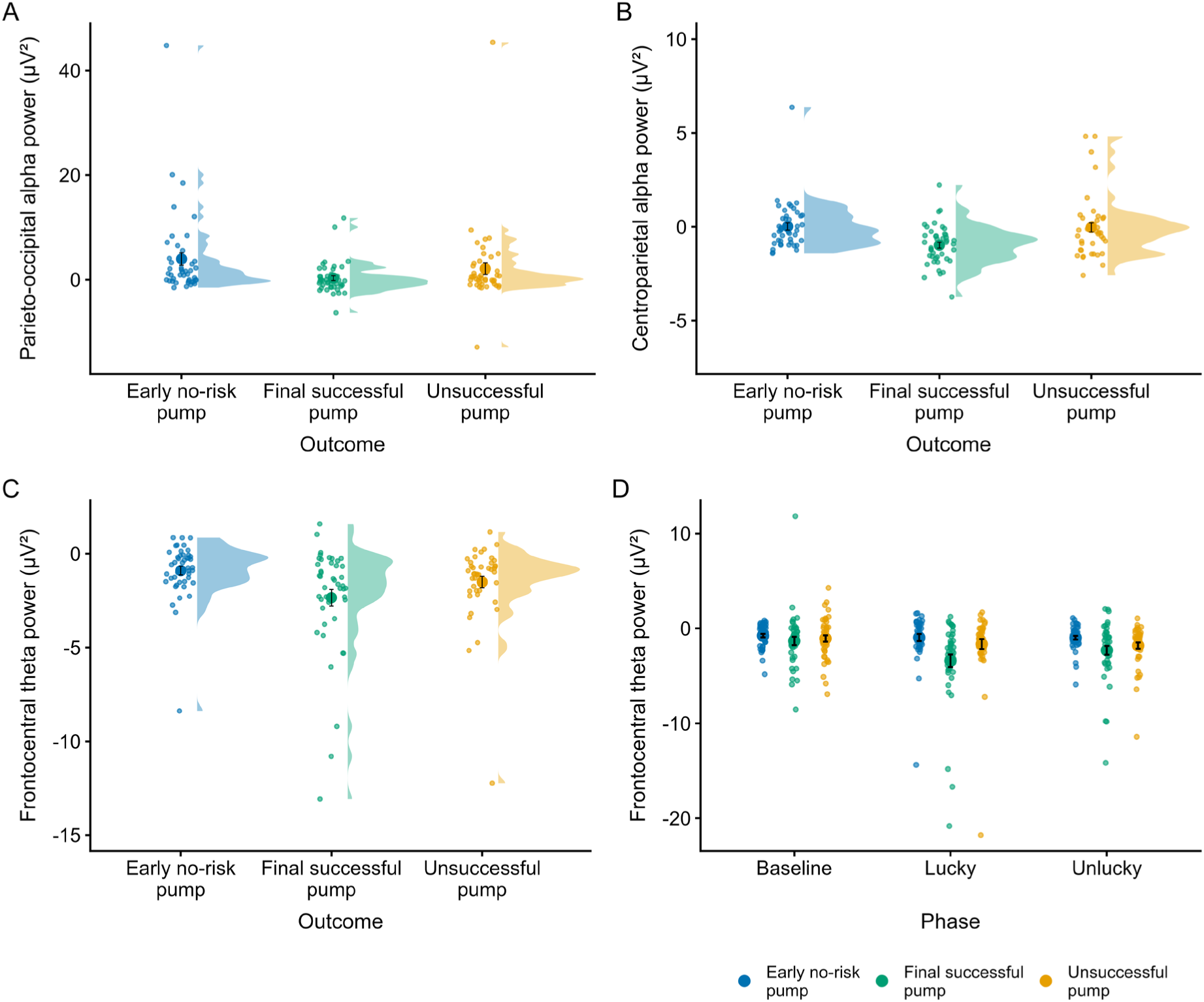
Total oscillatory power across outcome types and task phases. **(A)** Parieto-occipital alpha power split by outcome type, with the greatest alpha power increase after early no-risk pumps. **(B)** Centroparietal alpha power split by outcome type (early no-risk pump, final successful pump, unsuccessful pump), showing the strongest alpha suppression following final successful pumps. **(C)** Frontocentral theta power split by outcome types, demonstrating a graded pattern of theta suppression, most pronounced after final successful pumps. **(D)** Frontocentral theta power split by outcome type across the three task phases (baseline, lucky, unlucky), indicating that outcome-related differences in theta suppression were most prominent during the lucky and unlucky phases. Each panel displays individual data points as smaller circles (jittered to prevent overlap), group-level means as larger circles, and half-violin plots representing the distribution of power values. Error bars represent ±1 standard error of the mean (SEM).

To analyze basic behavioral data and mean total power, mixed-design and repeated-measures ANOVAs, respectively, were conducted. The factors involved in each analysis are specified below. The Greenhouse-Geisser epsilon correction (Greenhouse & Geisser, 1959) was used when necessary, i.e., when sphericity was violated as indicated by the significance of the Mauchly’s sphericity test. Original *df* values and corrected *p* values (if applicable) are reported together with partial eta-squared (*η^2^_p_*) as the measure of effect size. Multiple-comparison correction was applied within conceptually related families of analyses using Bonferroni correction. Because the primary analysis involved two behavioral ANOVAs and four time-frequency ANOVAs (see details below), behavioral and time**–**frequency ANOVAs were corrected separately, resulting in corrected significance thresholds of *p* < .025 (.05/2) and *p* < .0125 (.05/4), respectively. Supplementary analyses including Gender as an additional factor were treated as distinct families and corrected independently using the same procedure, resulting in corrected significance thresholds of *p* < .025 for behavioral ANOVAs and *p* < .0125 for time**–**frequency ANOVAs. The same Bonferroni-corrected significance threshold used in the main time**–**frequency analyses (*p* < .0125) was applied across all supplementary analyses testing for the robustness of time**–**frequency results to outlier removal. Post-hoc pairwise comparisons were performed using Tukey-adjusted tests. It should be noted that this paper does not offer a detailed analysis of behavioral data; for that analysis, please refer to the work of Bán et al. (2024).

## Results

### Behavioral results

The purpose of this analysis was to evaluate whether pumping behavior changed across the experimental phases differing in contextual uncertainty and to check the potential effect of order manipulation (i.e., the assignment of participants to Lucky-first or Unlucky-first order). We calculated each participants’ adjusted score (i.e., the mean number of pumps on unexploded balloons; Lejuez et al., 2002; Schmitz, Manske, Preckel, & Wilhelm, 2016) and their points collected in each phase. On both dependent variables separately, a mixed-design ANOVA was conducted with Order (Lucky-first vs. Unlucky-first) as a between-subjects factor and Phase (baseline vs. lucky vs. unlucky) as a within-subjects factor.

Considering the adjusted score, the main effect of Order showed no differences between the two Orders (*F*(1, 42) = 0.02, *p* = .886, *η*^2^*_p_* < .001), while the main effect of Phase revealed that the adjusted score differed in the three phases of the task (*F*(2, 84) = 235.04, *p* < .001, *η*^2^*_p_* = .85). Post-hoc analysis showed that the adjusted score was the highest in the lucky phase (*M* = 10.45), slightly lower in the baseline phase (*M* = 9.07) and the lowest in the unlucky phase (*M* = 4.59), as expected (all *p*s < .001). The Order * Phase interaction was not significant (*F*(2, 84) = 0.45, *p* = .591, *η*^2^*_p_* = .01).

Considering the points collected, the main effect of Order revealed comparable performance in the two Orders (*F*(1, 42) = 0.00, *p* = .970, *η*^2^*_p_* < .001), but the main effect of Phase showed that the points collected differed in the three phases (*F*(2, 84) = 194.73, *p* < .001, *η*^2^ = .82). Post-hoc analysis showed that participants achieved the highest score in the lucky phase (*M* = 3347.55), followed by the baseline phase (*M* = 2075.07), with the lowest scores observed in the unlucky phase (*M* = 494.30, all *p*s < .001). The Order * Phase interaction was not significant (*F*(2, 84) = 0.09, *p* = .826, *η*^2^*_p_* = .00).

These behavioral results as well as our previous work on learning rates (Bán et al., 2024) indicate that the order manipulation had no effect on task performance. Therefore, the Order factor was not included in the time–frequency analyses. To assess the potential influence of gender on the behavioral results, given well-documented gender differences in risk taking (Byrnes, Miller, & Schafer, 1999; Charness & Gneezy, 2012), including in BART performance (e.g., Cazzell, Li, Lin, Patel, & Liu, 2012; Lejuez et al., 2002; Lighthall, Mather, & Gorlick, 2009), we repeated these analyses including Gender as an additional between-subjects factor. Consistent with prior studies, men showed higher overall risk taking. Critically, however, Gender did not interact with either Phase or Order, nor with their interaction, suggesting that the effects of the experimental manipulations on behavior were comparable across genders. At the same time, given the relatively small and unbalanced male subsample, the absence of significant interactions should be interpreted with caution. Detailed statistics are reported in the Supplementary Materials (Tables S1 and S2).

### Time–frequency results

To analyze how reward anticipation evolved throughout the balloon inflation process in each phase, repeated-measures ANOVAs were conducted on the mean total power extracted for each frequency range. The ANOVA contained Outcome (early no-risk pump vs. final successful pump vs. unsuccessful pump) and Phase (baseline vs. lucky vs. unlucky) as within-subjects factors.

First, the increase of alpha power over the parieto-occipital region was analyzed. The main effect of Outcome (*F*(2, 86) = 7.86, *p* < .001, *η*^2^ = .16) showed a significant difference in oscillatory activity between the outcome types (Figure 2A and Figure 4A). Post-hoc analyses revealed that alpha power was higher following early no-risk pumps (*M* = 3.96 µV²) than after final successful pumps (*M* = 0.30 µV², *p* = .002), and showed a trend toward being higher compared to unsuccessful pumps (*M* = 2.11 µV², *p* = .091), while it was comparable between unsuccessful and final successful pumps (*p* = .136). The main effect of Phase (*F*(2, 86) = 0.37, *p* = .663, *η*^2^*_p_* = .01) and the Phase * Outcome interaction (*F*(4, 172) = 1.09, *p* = .353, *η*^2^*_p_* = .03) were not significant.

Second, an identical ANOVA was run for the decrease of alpha power at Cz. The main effect of Outcome (*F*(2, 86) = 16.78, *p* < .001, *η*^2^ = .28) revealed that oscillatory activity significantly differed between the outcome types (Figure 2B and Figure 4B). Post-hoc analysis showed that the decrease in alpha power was most prominent after final successful pumps (*M* = –0.98 µV², all *p*s < .001), but alpha power did not differ after unsuccessful pumps relative to early no-risk pumps (*M* = –0.04 µV² vs. *M* = 0.02 µV², *p* = .945). The main effect of Phase (*F*(2, 86) = 0.94, *p* = .389, *η*^2^ = .02) and the Phase * Outcome interaction (*F*(4, 172) = 1.03, *p* = .385, *η*^2^*_p_* = .02) were not significant.

Third, an identical ANOVA was conducted for the theta decrease at FCz. The main effect of Outcome (*F*(2, 86) = 14.57, *p* < .001, *η*^2^ = .25) revealed that oscillatory activity differed significantly between the outcome types. Post-hoc analysis showed a gradual change in theta power, with the most prominent decrease occurring after the final successful pumps (*M* = –2.35 µV²), followed by the unsuccessful pumps (*M* = –1.51 µV²), and then the early no-risk pumps (*M* = –0.90 µV², all *p*s ≤ .010, Figure 3 and Figure 4C). The main effect of Phase (*F*(2, 86) = 4.82, *p* = .022, *η*^2^*_p_* = .10) did not meet the corrected significance threshold (*p* < .0125). The Phase * Outcome interaction (*F*(4, 172) = 4.67, *p* = .003, *η*^2^*_p_* = .01, Figure 3 and Figure 4D) was also significant. Post-hoc analysis showed that the differences between the outcome types were most prominent in the lucky and unlucky phases. In the lucky phase, theta decrease was most pronounced following the final successful pumps (*M* = –3.41 µV², all *p*s < .002). The difference between the unsuccessful pumps (*M* = –1.66 µV²) and the early no-risk pumps (*M* = –0.96 µV²) reached trend level (*p* = .071). In the unlucky phase, oscillatory activity was comparable after the final successful pumps (*M* = –2.31 µV²) and the unsuccessful pumps (*M* = –1.81 µV², *p* = .328), while oscillatory activity was less negative after the early no-risk pumps (*M* = –0.98 µV²) compared to the other two outcome types (all *p*s < .001). In the baseline phase, none of the pairwise comparisons were significant (final successful pumps: *M* = –1.33 µV², unsuccessful pumps: *M* = –1.07 µV², early no-risk pumps: *M* = –0.76 µV², all *p*s > .339).

Fourth, an identical ANOVA was run for the beta power over the left centroparietal region. Neither the main effect of Outcome (*F*(2, 86) = 2.40, *p* = .109, *η*^2^*_p_* = .05), nor the main effect of Phase (*F*(2, 86) = 2.41, *p* = .099, *η*^2^*_p_* = .05), nor the Phase * Outcome interaction (*F*(4, 172) = 0.23, *p* = .895, *η*^2^*_p_* = .01) were significant.

To sum up the statistical findings, after the early no risk-pumps, a prominent increase in parieto-occipital alpha power was found. In addition, a pronounced decrease in centroparietal alpha power after the final successful pumps was also revealed. Over the frontocentral region, theta power showed a pronounced decrease following final successful pumps, with a graded pattern across outcomes. In addition, this effect was most prominent in the lucky and unlucky phases, while no differences between outcomes were observed in the baseline phase.

To test the potential influence of gender on the time–frequency results, we repeated these analyses on the mean total power including Gender as a between-subjects factor. In the alpha and beta bands, no significant main effects or interactions involving Gender were observed. In contrast, for theta activity, significant main and lower-order interaction effects involving Gender were observed. However, the three-way interaction (Gender * Outcome * Phase) was not significant, suggesting that the primary Outcome * Phase effect was broadly comparable across genders. At the same time, given the relatively small and unbalanced male subsample, the present data do not allow for a conclusive interpretation of the non-significant higher-order interaction. Detailed statistics are reported in the Supplementary Materials (Table S3).

Moreover, we conducted additional robustness analyses to assess the potential influence of outliers. Outliers were identified in the power values, and all analyses were repeated on samples excluding these observations (i.e., restricted samples). The pattern of results remained largely similar compared to the full sample, indicating that the findings are robust to the exclusion of extreme values. Detailed statistics are reported in the Supplementary Materials (Table S4).

To further explore the decrease in theta-band activity and relate the theta suppression observed particularly after the final successful pumps to behavioral indices, we conducted two correlational analyses. First, we tested whether theta decrease was associated with reaction times (RTs) for the cash-out decisions following the final successful pumps. Second, we tested whether theta decrease was related to RTs for the final successful pumps. While no significant association was observed between theta decrease and RTs for cash-out decisions, RTs for the final successful pumps positively correlated with theta decrease (*r_s_*(40) = 0.434, *p* = .004). Because more negative theta values indicate stronger suppression, this association suggests that faster execution of the final successful pump was accompanied by stronger post-response theta suppression. Detailed statistics and further information are reported in the Supplementary Materials (Figure S1).

## Discussion

The present study investigated the neurocognitive mechanisms underlying reward anticipation during sequential decision making. To this end, EEG oscillatory power was measured time-locked to the critical responses of the decision-making process implemented in a modified BART. In this ambiguity-based decision environment, reward probability changed across the task according to predetermined probability functions that were unknown to participants. In terms of the EEG results, two distinct patterns emerged in the alpha frequency range. Right before the reward (inflated balloon), parieto-occipital alpha power was increased at the early no-risk inflation step compared to both final successful and unsuccessful inflations steps. Meanwhile, alpha power was suppressed over centroparietal electrode sites for a more extended time after the final successful balloon inflation. This centroparietal alpha suppression was mostly lacking after the unsuccessful balloon inflation and the early no-risk inflation step. Furthermore, frontocentral theta power showed a pronounced decrease after the final successful balloon inflation, especially in the lucky phase. Together, these results indicate that EEG spectral components are sensitive to within-trial escalating risk, even in the absence of explicit predictive cues.

Behavioral results clearly demonstrated that participants adapted to the different task phases (baseline, lucky, unlucky), indicating sensitivity to the broader, long-term probability structure of the task. In contrast, oscillatory activity during anticipation was more strongly modulated by the immediate, within-trial escalation of risk than by contextual uncertainty. Although contextual uncertainty effects were not entirely absent at the EEG level, their influence was comparatively weaker than that of inflation-step-specific risk dynamics.

This pattern suggests a dissociation across levels of processing. Behavioral indices accumulate information across trials and therefore reflect integration of the broader probability structure characterizing each phase, consistent with prior work demonstrating rapid adaptation to probabilistic reward regularities in the BART (Bonini et al., 2018; Kardos et al., 2016; Kóbor et al., 2021; Kóbor et al., 2023; Koscielniak et al., 2016; Petropoulos Petalas et al., 2020; Smith et al., 2016; Young & McCoy, 2019). This is also consistent with our earlier findings showing that contextual uncertainty manipulations shape decision making (Bán et al., 2024). By contrast, anticipatory neural activity in the analyzed time window appeared to be more tightly coupled to the dynamically increasing probability of loss associated with the current inflation step. Thus, while participants adjusted their overall behavioral strategy to contextual uncertainty, the momentary anticipatory brain state was primarily driven by the evolving hazard within each balloon. Within this framework, the apparent dissociation does not indicate inconsistency between behavioral and EEG measures but rather suggests that they index complementary aspects of adaptive processing operating at different levels of the task structure.

Oscillatory power modulations do not follow a fixed “positive” or “negative” activation pattern across frequency bands; rather, both increases and decreases in power may reflect different functional states depending on task demands. In the case of alpha oscillations, studies investigating the spectral correlates of cognitive mechanism have found that alpha power increases when cognitive effort is reduced and decreases when attention and task engagement are intensified. This phenomenon is thought to reflect functional inhibition and cortical activation, respectively (e.g., Bastiaansen & Brunia, 2001; Glazer et al., 2018; Klimesch, 2012). In line with this view and our hypothesis, the increase in parieto-occipital alpha power at the early no-risk inflation step before presenting a sure reward suggests a disengagement from active anticipation. This synchronized alpha band rhythm can be considered as the operation of a “resting” or “idling” cognitive state (Pfurtscheller, Stancák, & Neuper, 1996). At this early no-risk step, effortlessly inflating the balloon further while maintaining a steady cognitive state was critical instead of expecting a certain outcome. Thus, weighing the odds of a balloon burst should have been inhibited to proceed with the task to accumulate higher earnings. Despite the absence of explicit information regarding the zero probability of the balloon bursting after the initial two pumps, it is reasonable to assume that the participants deduced this information quickly upon completing some balloons. This assumption is further substantiated by the ecological implausibility of a very early burst, given the inherent features of real balloons. Importantly, this increase in alpha power was diminished after the unsuccessful inflation step and was virtually absent after the final successful one. The latter finding suggests that neurocognitive mechanisms different from “idling” were engaged at the late stage of the sequential decision-making process, which we discuss next.

While we found parieto-occipital alpha increase at the early no-risk inflation step, a distinct centroparietal alpha suppression emerged following the final successful balloon inflation. This effect was selective to this late stage and was not observed after unsuccessful or early no-risk pumps. This suppression likely reflects increased anticipatory attention and outcome-related expectancy in the post-response interval (Glazer et al., 2018; Pornpattananangkul & Nusslock, 2016; van den Berg et al., 2014). At this high-risk inflation step, participants might have engaged in evaluating the consequences of their action and may have prepared to collect the accumulated earnings, despite being unaware of the actual outcome of their pump (Helfinstein et al., 2014). In contrast, unsuccessful pumps did not show comparable suppression but rather a slight parieto-occipital alpha increase, suggesting that participants may have maintained a task-oriented state geared toward continued pumping, similar to the early no-risk inflation step. This dissociation between successful and unsuccessful pumps suggests that substantially different outcomes are anticipated at the end of these balloon trials (Helfinstein et al., 2014; Kardos et al., 2016). Furthermore, the absence of centroparietal alpha suppression at the early no-risk inflation step implies that attentional and expectancy processes were minimally engaged when participants did not need to anticipate potential rewards or losses.

The centroparietal distribution of the alpha suppression was not expected based on prior studies, which have more commonly reported decreases in alpha power over occipital and parieto-occipital sites (Bastiaansen et al., 2002; Bastiaansen & Brunia, 2001; Gómez et al., 2004; Pornpattananangkul & Nusslock, 2016; van den Berg et al., 2014; Zhang et al., 2023). However, previous work has also reported more anterior alpha effects associated with task-relevant processing. For example, van den Berg et al. (2014) found that not only occipital alpha but also frontocentral alpha differentiated reward-prospect and no-reward-prospect trials of a cognitive conflict task, considered as the marker of increased preparatory attention to suppress irrelevant information. Moreover, the frequency range of the present effect may overlap with lower alpha activity (approximately 6–10 Hz), which has been linked to expectancy-related and attentional processes and is known to exhibit a broader, more anterior topography (Gómez et al., 2004; Klimesch, 1999; Klimesch, Pfurtscheller, Mohl, & Schimke, 1990). Although our analyses did not explicitly distinguish between lower and upper alpha sub-bands, the centroparietal alpha suppression observed following final successful pumps likely reflects a combination of heightened attentional engagement and reward-related expectancy in the post-response interval preceding outcome realization.

Contrary to our initial hypothesis, final successful balloon inflations were accompanied by decreases in both centroparietal alpha and frontocentral theta power. We had tentatively predicted reduced theta at the early no-risk step relative to the final inflation steps, based on prior reports linking higher reward probability to lower theta power (Bunzeck et al., 2011; Doñamayor et al., 2012). Instead, theta power was lowest following the final successful pump. Although this pattern differs from studies reporting increased anticipatory theta before potential rewards (Alí Diez et al., 2024; Reinhart & Woodman, 2014; Zhang et al., 2023), the literature on anticipatory theta is mixed. Importantly, midfrontal theta has been widely associated with conflict monitoring, performance monitoring, uncertainty processing, and the need for cognitive control, rather than reward probability per se (Botvinick, Cohen, & Carter, 2004; Cavanagh & Frank, 2014; Cohen & Donner, 2013). From this perspective, the observed theta decrease may reflect reduced cognitive control or conflict monitoring demands following commitment to the selected pump action. In the present response-locked, self-paced design, execution of the final successful pump may mark stabilization of the chosen action, reducing the need for continued control engagement despite ongoing outcome uncertainty, and thereby attenuating theta activity in line with control-related interpretations of midfrontal theta dynamics (Cavanagh, Cohen, & Allen, 2009). Consistent with this interpretation, the observed brain–behavior correlation indicates that faster responses on the final successful pump were followed by stronger theta suppression in the subsequent post-response interval.

Differences in task structure may help explain the discrepancy with prior findings. Studies reporting anticipatory theta increases have employed externally cued paradigms in which reward probability is explicitly signaled and anticipation is stimulus locked. In such contexts, theta modulation may reflect preparation for action or evaluation under externally imposed uncertainty. In contrast, the BART involves sequential, self-paced decisions in which reward expectancy evolves gradually and is inferred from experience rather than cued explicitly. Moreover, our analyses were response-locked, capturing neural dynamics following an active decision rather than in response to an external cue. Under these conditions, anticipatory theta may primarily index internal decision resolution and dynamic regulation of cognitive control rather than explicit encoding of reward probability. This would also be in line with the general theta suppression observed in all conditions, indicating action commitment throughout the balloon inflation process. Additionally, because the final successful pump is followed by trial termination (i.e., cash-out response), more reduced theta may partly reflect diminished need for continued action preparation or monitoring compared to preceding pumps (but see Helfinstein et al., 2014).

The phase-wise modulation further informs this interpretation. Theta decreases across within-trial risk levels were most pronounced in the lucky phase, where the burst probability function was flatter and extended reward pursuit was possible. In this context, successful pumps may have been associated with stronger internal confirmation of decision adequacy and reduced monitoring demands. By contrast, in the unlucky phase, elevated loss probability may have sustained cognitive control engagement, attenuating the theta decrease. Thus, frontocentral theta appears sensitive not simply to contextual uncertainty, but to the balance between control demands and internally inferred decision confidence across escalating within-trial risk and varying reward–loss contingencies.

It is also important to consider that balloon size, which increased across inflation steps, may have served as an intrinsic reward-magnitude cue and influenced visual salience or attentional processing. Larger balloons at later inflation steps may therefore have elicited stronger visual or attentional responses independent of probabilistic reward expectations. Although this represents a potential perceptual confound, centroparietal alpha and frontocentral theta activity differentiated between procedurally similar final balloon inflations depending on their forthcoming, yet unrevealed, outcome. This indicates that anticipatory neural dynamics were not solely driven by visual magnitude cues but were also sensitive to learned expectations about the probabilistic reward structure (Bonini et al., 2018; Helfinstein et al., 2014; Kardos et al., 2016; Koscielniak et al., 2016; Petropoulos Petalas et al., 2020). Moreover, because final successful and unsuccessful pumps could occur at any inflation step, they were not associated with fixed visual magnitudes. Future studies, nevertheless, should more directly disentangle perceptual magnitude from probabilistic expectancy and clarify how anticipatory theta dynamics are shaped by task structure, temporal locking (stimulus- vs. response-locked analyses), and whether anticipation is externally cued or internally generated. Given that substantially more findings are available regarding theta modulation at the reward-outcome stage (Glazer et al., 2018), such investigations will be essential for establishing the functional role of theta oscillations during reward anticipation. Overall, the present findings suggest that, in sequential and self-paced risk-taking contexts, anticipatory theta dynamics may primarily reflect adaptive regulation of cognitive control and commitment to the selected action rather than straightforward encoding of reward probability.

Although we formulated a priori hypotheses regarding beta power modulations – based on prior findings linking anterior beta increases to reward anticipation and incentive motivation (Apitz & Bunzeck, 2014; Bunzeck et al., 2011; Reinhart & Woodman, 2014; Xiao et al., 2024) – we did not observe reliable beta modulation as a function of escalating risk or contextual uncertainty. This absence of modulation was evident both in the time–frequency inspection and in the region-of-interest analysis of total power. The lack of beta effects therefore warrants careful consideration.

Our prediction assumed that escalating reward probability within the BART would parametrically increase incentive salience and thereby enhance anticipatory motor readiness. However, the motivational structure of the BART differs from paradigms in which beta increases have been most consistently reported. In many cue-based reward tasks, high versus low incentives are explicitly signaled and temporally isolated, producing relatively unambiguous motivational states. In contrast, the BART involves a gradual and self-paced escalation of risk in which increasing potential gains are coupled with increasing burst probability. Thus, reward magnitude and hazard co-escalate, generating a mixed motivational signal that simultaneously supports approach and caution. Under these circumstances, incentive-related beta dynamics may not manifest as clear monotonic or condition-specific increases during anticipation.

Moreover, beta oscillations have been proposed to index maintenance of the current sensorimotor or cognitive set rather than incentive value per se (Engel & Fries, 2010). From this perspective, beta synchronization stabilizes the status quo and supports ongoing motor states. In the present task, participants repeatedly executed the same inflation response across risk levels, and the decision to cash out did not introduce a qualitatively distinct motor program until commitment. Consistent with work emphasizing beta’s role in motor control and movement stability (Jenkinson & Brown, 2011), the comparable beta activity observed across experimental conditions may therefore reflect sustained maintenance of the ongoing response set while awaiting outcome resolution, rather than modulation by reward probability.

Notably, whereas alpha and theta activity differentiated risk levels prior to outcome, beta activity remained insensitive to escalating hazard, suggesting functional dissociation rather than a unitary anticipatory mechanism. Although increased beta has also been linked to statistical learning (Bogaerts, Richter, Landau, & Frost, 2020), the present findings indicate that learned reward regularities embedded within a sequential risk structure do not necessarily translate into differential beta modulation during anticipation. Taken together, the null beta result helps refine the interpretation of beta oscillations in reward contexts. It suggests that beta power may be particularly sensitive to explicitly cued incentive manipulations and clear shifts in motor preparation, whereas gradually escalating, covertly structured risk – as implemented in the BART – may preferentially engage other oscillatory mechanisms.

While our results advance the understanding of reward anticipation in sequential decision making, some methodological limitations should be considered. Focusing only on the anticipation phase in a response-locked manner allowed us to capture neurocognitive responses that were relatively comparable across phases and inflation steps. In this task version, analyzing the outcome stage in a stimulus-locked manner involves comparing either different visual stimuli for rewarded and non-rewarded trials (i.e., inflated balloon or cash-out screen vs. balloon burst; see e.g., Kóbor et al., 2015) or various reward magnitudes (balloon sizes) if only the successfully inflated balloons are considered (Kardos et al., 2016). Even in the present study, the short duration of the anticipatory window, the challenge of establishing an EEG segment baseline without stimulus presentation and response preparation, and the potentially confounding balloon size as a reward cue should be considered in interpreting these results. For instance, the short anticipatory window did not allow us to reliably measure delta oscillations that have been found to be modulated mostly in the cue-evaluation substage of reward anticipation (Glazer et al., 2018). Nonetheless several of these limitations stem from characteristics of the paradigm, these features were necessary to preserve ecological validity and capture the dynamics of sequential risk taking.

Although gender differences in overall risk taking were observed at the behavioral level, and differences in theta activity were observed in lower-order effects, gender did not significantly interact with the critical effects of the experimental manipulations on either behavior or oscillatory dynamics. Specifically, the Outcome by Phase interaction – reflecting within-trial modulation of reward anticipation in theta activity – was broadly comparable across genders. This pattern suggests that gender-related differences might be more evident at the level of baseline behavior and neural activity, whereas the dynamic adjustment of anticipatory processes to changing risk and uncertainty might be relatively similar across groups. At the same time, the limited number of male participants reduces the statistical power to detect potential gender-related interactions, warranting caution in interpreting the non-significant higher-order effects. Future studies with more balanced samples are needed to more conclusively assess potential gender-related modulation.

Despite the constraints, the present findings advance our understanding of reward anticipation by demonstrating that oscillatory activity differentiates within-trial risk levels prior to feedback, even in the absence of explicit reward cues. Parieto-occipital alpha increases at early no-risk pumps suggest relatively reduced deliberative engagement when reward is certain, whereas centroparietal alpha and frontocentral theta suppressions at later inflation steps indicate heightened attentional engagement and adaptive regulation of cognitive control following commitment to the selected action. Collectively, these results suggest that distinct neurocognitive mechanisms are engaged at different stages of the sequential decision-making process as conditional risk unfolds. Future research should further clarify how task structure and temporal dynamics shape alpha and theta oscillations during reward anticipation and adaptive decision making.

## Declarations

### Ethics approval and consent to participate

The study was conducted in accordance with the Declaration of Helsinki. The United Ethical Review Committee for Research in Psychology (EPKEB) in Hungary gave approval to the procedure. Written informed consent was obtained from all participants.

### Consent for publication

Not applicable.

### Data availability statement

Data supporting the findings of this study (processed behavioral and time–frequency data) are available via the Open Science Framework (https://osf.io/bwzfa/files/osfstorage), located in the BART luck FC TF folder. EEG data in various formats (raw, pre-processed, segmented, etc.) will be provided from the corresponding author (A.K.) upon written request. The experiment was not preregistered.

### Code availability statement

Not applicable as no custom code was used to process the data described in the study.

### Competing interests

The authors declare no competing interests.

### Funding information

This research was supported by project no. 124412 (PI: A.K.) that have been implemented with the support provided by the Ministry of Innovation and Technology of Hungary from the National Research, Development and Innovation Fund, financed under the FK funding scheme. Furthermore, the research was supported by the National Brain Research Program by the Hungarian Academy of Sciences (project NAP2022-I-1/2022) and the János Bolyai Research Scholarship of the Hungarian Academy of Sciences.

## Acknowledgements

The authors thank Noémi Éltető and Zsófia Kardos for their helpful suggestions on the experimental design. They also thank the help of Dóra Bárány in EEG data acquisition, and the useful comments of Dezső Nemeth and Ferenc Honbolygó on the earlier version of the manuscript.

## CRediT author statement

**Eszter Tóth-Fáber:** Conceptualization, Methodology, Project administration, Investigation, Formal analysis, Data Curation, Writing – Original Draft, Writing – Review & Editing, Visualization.

**Andrea Kóbor:** Conceptualization, Methodology, Software, Project administration, Formal analysis, Data Curation, Supervision, Funding acquisition, Writing – Original Draft, Writing – Review & Editing.

## Use of AI Generated Content (AIGC) and tools

During the preparation of this manuscript, the authors used the AI-based tool ChatGPT (OpenAI) for language editing and stylistic refinement of the English text. The tool was not used for data analysis or to generate scientific conclusions. All outputs were critically reviewed and edited by the authors, who take full responsibility for the content of the publication.

## Notes

https://osf.io/3pczq

### Competing Interest Statement

The authors have declared no competing interest.

### Summary of Updates

We revised the presentation of the supplementary analyses involving gender and now interpret the results in a more cautious manner. Moreover, we explored brain-behavior correlations in terms of theta suppression following final successful pumps. We also employed Bonferroni-correction throughout the analyses to control for multiple comparisons.

