## Supplementary Materials for "Alpha and theta oscillations differentiate escalating risk levels during reward anticipation in sequential decision making"

### Justification of sample size

The target sample size was determined based on prior EEG investigations of reward anticipation as well as practical considerations inherent to time–frequency EEG data collection. A total of 50 participants were initially tested (25 per order condition: Lucky-first vs. Unlucky-first). Following predefined artifact-rejection procedures, six participants were excluded (cf. EEG recording and analyses section in the main text). The final sample therefore consisted of 44 participants.

The final sample size is comparable to or larger than that of many prior EEG time–frequency studies examining reward anticipation and related processes, which commonly include samples in the range of approximately 20–40 participants (e.g., Doñamayor, Schoenfeld, & Münte, 2012; Pornpattananangkul & Nusslock, 2016; van den Berg, Krebs, Lorist, & Woldorff, 2014; Zhang et al., 2023). It is also similar to or larger than several previous EEG investigations of risk taking in the BART (e.g., Gu, Zhang, Luo, Wang, & Broster, 2018; Kardos et al., 2016; Kóbor et al., 2015) and comparable to recent time–frequency BART studies (e.g., Qianlan, Shou, Tianya, Wei, & Liu, 2025; Tao et al., 2023).

In addition, the study employed a fully within-subject 3 (Outcome: early no-risk pump vs. final successful pump vs. unsuccessful pump) × 3 (Phase: baseline vs. lucky vs. unlucky) repeated-measures design. Estimating the effects of interest within participants reduces inter-individual variability in the critical contrasts and increases statistical sensitivity.

Furthermore, EEG data quality was high. The mean proportion of removed segments was low and comparable across Outcome levels (early no-risk: 6.29%, unsuccessful: 5.35%, final successful: 4.43%, cf. main text), indicating that artifact rejection did not differentially affect specific conditions. Importantly, the time–frequency analyses were based on a relatively high number of artifact-free segments per condition. On average, participants contributed approximately 30–75 artifact-free segments per Outcome level across the phases (cf. main text). Even the lowest-frequency cells retained more than 10 artifact-free segments per participant.

This trial density supports stable spectral power estimation at the single-subject level and reliable group-level comparisons.

In summary, although no formal a priori power analysis was conducted, the combination of a repeated-measures factorial design, low and uniform artifact rejection rates, and substantial per-condition trial counts provides sufficient sensitivity to detect effects commonly reported in similar EEG paradigms.

### Testing the possible influence of gender on the behavioral and EEG results

To examine whether the observed gender imbalance influenced the pattern of effects, we conducted additional analyses including Gender as a between-subjects factor in the ANOVAs. Multiple-comparison correction was applied within conceptually related families of analyses using Bonferroni correction. Hence, behavioral analyses including gender were evaluated against a corrected threshold of $p < .025$, whereas time–frequency analyses including gender were evaluated against a corrected threshold of $p < .0125$.

For the behavioral analyses, we observed overall gender differences in risk taking; however, Gender did not interact with any of the experimental factors (Order, Phase, or their interaction). For the EEG analyses, no significant main effects or interactions involving Gender were observed in the alpha and beta bands. For theta activity, significant main and lower-order interaction effects involving Gender were observed, indicating overall and condition-specific differences in theta dynamics between men and women. Critically, however, the three-way interaction (Gender * Outcome * Phase) was not significant, indicating that the primary Outcome * Phase effect did not differ by gender. Altogether, these analyses indicate that the reported experimental effects are unlikely to be driven by gender differences. However, given the relatively small number of male participants, these analyses may have limited power to detect moderation effects; thus, the absence of interaction should be interpreted cautiously. The exact statistics are presented in Tables S1, S2, and S3.

**Table S1.** The effect of gender on the mean adjusted number of pumps.

| Factors | Statistics |
|---|---|
| Gender | **$F(1, 40) = 5.66, p = .022, \eta^2_p = .12$** |
| Order | $F(1, 40) = 0.02, p = .884, \eta^2_p < .001$ |
| Phase | $F(2, 80) = 147.26, p < .001, \eta^2_p = .79$ |
| Gender * Order | $F(1, 40) = 0.57, p = .456, \eta^2_p = .01$ |
| Gender * Phase | $F(2, 80) = 1.33, p = .267, \eta^2_p = .03$ |
| Order * Phase | $F(2, 80) = 0.06, p = .896, \eta^2_p = .00$ |
| Gender * Order * Phase | $F(2, 80) = 0.07, p = .892, \eta^2_p = .00$ |

*Notes.* All significant ($p < .025$) main effects and interactions involving Gender are presented in **bold**. The main effect of Gender revealed that men and women differed in risk taking. Post-hoc analysis showed that men had higher adjusted score ($M = 9.21$) than women ($M = 7.75$). The main effect of Phase revealed that the adjusted score differed in the three phases of the task. Post-hoc analysis showed that the adjusted score was the highest in the lucky phase ($M = 10.98$), slightly lower in the baseline phase

($M$ = 9.66), and the lowest in the unlucky phase ($M$ = 4.80), as expected (all $p$s < .004). Importantly, influence of gender on task manipulation was not detected.

**Table S2.** The effect of gender on total points collected in the task.

| Factors | Statistics |
|---|---|
| Gender | $F(1, 40) = 2.38, p = .130, \eta^2_p = .06$ |
| Order | $F(1, 40) = 0.03, p = .855, \eta^2_p < .001$ |
| Phase | $F(2, 80) = 126.17, p < .001, \eta^2_p = .76$ |
| Gender * Order | $F(1, 40) = 0.01, p = .911, \eta^2_p < .001$ |
| Gender * Phase | $F(2, 80) = 1.51, p = .229, \eta^2_p = .04$ |
| Order * Phase | $F(2, 80) = 0.01, p = .941, \eta^2_p < .001$ |
| Gender * Order * Phase | $F(2, 80) = 0.22, p = .697, \eta^2_p = .01$ |

*Notes.* The main effect of Phase showed that the points collected differed in the three phases. Post-hoc analysis showed that participants achieved the highest score in the lucky phase ($M$ = 3542), followed by the baseline phase ($M$ = 2154), with the lowest scores observed in the unlucky phase ($M$ = 476, all $p$s < .001).

**Table S3.** The effect of gender on the total power in the theta, alpha, and beta frequency bands.

| Factors | Statistics |
|---|---|
| Alpha increase | |
| Gender | $F(1, 42) = 0.46, p = .503, \eta^2_p = .01$ |
| Phase | $F(2, 84) = 0.09, p = .882, \eta^2_p = .00$ |
| Outcome | $F(2, 84) = 5.57, p = .006, \eta^2_p = .12$ |
| Gender * Phase | $F(2, 84) = 1.39, p = .255, \eta^2_p = .03$ |
| Gender * Outcome | $F(2, 84) = 0.37, p = .683, \eta^2_p = .01$ |
| Phase * Outcome | $F(4, 168) = 0.08, p = .966, \eta^2_p = .00$ |
| Gender * Phase * Outcome | $F(4, 168) = 1.23, p = .302, \eta^2_p = .03$ |
| Alpha decrease | |
| Gender | $F(1, 42) = 3.54, p = .067, \eta^2_p = .08$ |
| Phase | $F(2, 84) = 0.40, p = .656, \eta^2_p = .01$ |
| Outcome | $F(2, 84) = 10.29, p < .001, \eta^2_p = .20$ |
| Gender * Phase | $F(2, 84) = 0.05, p = .941, \eta^2_p = .00$ |
| Gender * Outcome | $F(2, 84) = 0.02, p = .963, \eta^2_p < .001$ |
| Phase * Outcome | $F(4, 168) = 0.68, p = .577, \eta^2_p = .02$ |
| Gender * Phase * Outcome | $F(4, 168) = 0.54, p = .663, \eta^2_p = .01$ |

| | |
|---|---|
| Theta decrease | |
| Gender | **$F(1, 42) = 15.23, p < .001, \eta^2_p = .27$** |
| Phase | $F(2, 84) = 8.59, p = .002, \eta^2_p = .17$ |
| Outcome | $F(2, 84) = 25.45, p < .001, \eta^2_p = .38$ |
| Gender * Phase | $F(2, 84) = 3.72, p = .045, \eta^2_p = .08$ |
| Gender * Outcome | **$F(2, 84) = 8.69, p = .002, \eta^2_p = .17$** |
| Phase * Outcome | $F(4, 168) = 5.49, p < .001, \eta^2_p = .12$ |
| Gender * Phase * Outcome | $F(4, 168) = 1.42, p = .237, \eta^2_p = .03$ |
| Beta increase | |
| Gender | $F(1, 42) = 1.47, p = .232, \eta^2_p = .03$ |
| Phase | $F(2, 84) = 0.87, p = .419, \eta^2_p = .02$ |
| Outcome | $F(2, 84) = 1.72, p = .193, \eta^2_p = .04$ |
| Gender * Phase | $F(2, 84) = 1.55, p = .220, \eta^2_p = .04$ |
| Gender * Outcome | $F(2, 84) = 0.67, p = .478, \eta^2_p = .02$ |
| Phase * Outcome | $F(4, 168) = 0.33, p = .829, \eta^2_p = .01$ |
| Gender * Phase * Outcome | $F(4, 168) = 0.92, p = .442, \eta^2_p = .02$ |

*Notes*. All significant ($p < .0125$) main effects and interactions involving Gender are highlighted in **bold**. For theta decrease, a significant main effect of Gender was observed, indicating an overall difference between men and women. Post-hoc analyses showed that men had a more negative activation ($M = -3.73$ µV²) than women ($M = -1.11$ µV²). The Gender * Phase interaction did not meet the corrected significance threshold ($p < .0125$). The Gender * Outcome interaction reached significance. In men, post-hoc analyses revealed that all outcome types differed significantly: the greatest decrease occurred after the final successful pump ($M = -5.73$ µV²), followed by the unsuccessful pump ($M = -3.27$ µV²) and the early no-risk pump ($M = -2.19$ µV², all *p*s < .025). In women, the pattern was similar, with the largest decrease after the final successful pump ($M = -1.60$ µV²), which was significantly more negative than after the early no-risk pump ($M = -0.62$ µV², $p = .016$), but did not differ from the unsuccessful pump ($M = -1.12$ µV², $p = .204$). The early no-risk vs. unsuccessful pump comparison was significant ($p = .026$).

### Robustness to outlier removal

We tested the robustness of our time–frequency results to outlier removal. First, it is important to note that from the analyses presented in the manuscript, participants showing insufficient EEG data quality (i.e., low segment numbers due to noisy data, $n = 6$) were already excluded.

We conducted outlier detection on the power values and we additionally repeated all analyses on restricted samples. We implemented both strict and lenient outlier exclusion criteria. Regarding both, participants were defined as extreme outliers as data points exceeding 3 × IQR below the first quartile (Q1) or above the third quartile (Q3). Under the strict criterion, participants were excluded if they were identified as extreme outliers in any condition. Under the lenient criterion, participants were excluded only if they were classified as extreme outliers in more than half of the conditions (i.e., at least in five conditions out of nine conditions). The results for the two restricted samples, alongside the results on the full sample, are presented in Table S4. As these analyses represented robustness checks within the same inferential family, the same Bonferroni-corrected significance threshold used in the main analyses ($p < .0125$) was applied across all analyses. Notably, the majority of findings were consistent with those obtained from the full sample, indicating that outliers did not confound the original results.

**Table S4.** Time–frequency results on the whole sample as well as restricted, outlier-filtered samples.

| | Full sample | Lenient exclusion | Strict exclusion |
|---|---|---|---|
| | Alpha increase | | |
| N | 44 | 43 | 39 |
| Outcome | **$F(2, 86) = 7.86$, $p < .001$, $\eta^2_p = .16$** | **$F(2, 84) = 8.45$, $p = .001$, $\eta^2_p = .17$** | **$F(2, 76) = 14.67$, $p < .001$, $\eta^2_p = .28$** |
| Phase | $F(2, 86) = 0.37$, $p = .663$, $\eta^2_p = .01$ | $F(2, 84) = 1.17$, $p = .306$, $\eta^2_p = .03$ | $F(2, 76) = 3.41$, $p = .054$, $\eta^2_p = .08$ |
| Phase * Outcome | $F(4, 172) = 1.09$, $p = .353$, $\eta^2_p = .03$ | $F(4, 168) = 1.56$, $p = .210$, $\eta^2_p = .04$ | $F(4, 152) = 1.38$, $p = .255$, $\eta^2_p = .04$ |
| | Alpha decrease | | |
| N | 44 | 43 | 38 |
| Outcome | **$F(2, 86) = 16.78$, $p < .001$, $\eta^2_p = .28$** | **$F(2, 84) = 15.30$, $p < .001$, $\eta^2_p = .27$** | **$F(2, 74) = 20.77$, $p < .001$, $\eta^2_p = .36$** |

| | | | |
|---|---|---|---|
| Phase | $F(2, 86) = 0.94$, $p = .389$, $\eta^2_p = .02$ | $F(2, 84) = 0.89$, $p = .402$, $\eta^2_p = .02$ | $F(2, 74) = 0.04$, $p = .936$, $\eta^2_p < .001$ |
| Phase * Outcome | $F(4, 172) = 1.03$, $p = .385$, $\eta^2_p = .02$ | $F(4, 168) = 0.90$, $p = .444$, $\eta^2_p = .02$ | $F(4, 148) = 1.23$, $p = .303$, $\eta^2_p = .03$ |
| | | Theta decrease | |
| N | 44 | 43 | 40 |
| Outcome | **$F(2, 86) = 14.57$, $p < .001$, $\eta^2_p = .25$** | **$F(2, 84) = 13.29$, $p < .001$, $\eta^2_p = .24$** | **$F(2, 78) = 12.33$, $p < .001$, $\eta^2_p = .24$** |
| Phase | $F(2, 86) = 4.82$, $p = .022$, $\eta^2_p = .10$ | $F(2, 84) = 4.66$, $p = .018$, $\eta^2_p = .10$ | $F(2, 78) = 2.96$, $p = .066$, $\eta^2_p = .07$ |
| Phase * Outcome | **$F(4, 172) = 4.67$, $p = .003$, $\eta^2_p = .01$** | **$F(4, 168) = 4.65$, $p = .003$, $\eta^2_p = .10$** | $F(4, 156) = 2.79$, $p = .037$, $\eta^2_p = .07$ |
| | | Beta increase | |
| N | 44 | | 43 |
| Outcome | $F(2, 86) = 2.40$, $p = .109$, $\eta^2_p = .05$ | | $F(2, 84) = 1.58$, $p = .216$, $\eta^2_p = .04$ |
| Phase | $F(2, 86) = 2.41$, $p = .099$, $\eta^2_p = .05$ | | $F(2, 84) = 4.39$, $p = .016$, $\eta^2_p = .10$ |
| Phase * Outcome | $F(4, 172) = 0.23$, $p = .895$, $\eta^2_p = .01$ | | $F(4, 168) = 0.43$, $p = .746$, $\eta^2_p = .01$ |

*Notes*. All significant ($p < .0125$) main effects and interactions are highlighted in **bold**. The lenient exclusion column for beta increase is empty because there were no participants who were outliers in more than half of the conditions. The results obtained from the lenient and strict samples were largely consistent with those from the full sample, indicating that outlier handling did not meaningfully affect the main findings. Slight differences in the post-hoc results were found regarding the decrease in theta power. For the main effect of Outcome, post-hoc results were identical across all three samples. The interaction did not meet the corrected significance level in the strict sample. Across the full and lenient samples, post-hoc patterns were identical.

### Brain-behavior associations

To further explore the decrease in theta-band activity and relate the theta suppression observed particularly after the final successful pump to behavioral indices, we conducted two correlational analyses. Specifically, we examined associations between theta decrease following the final successful pumps and two behavioral measures: (1) reaction times (RTs) for the cash-out decisions following the final successful pumps and (2) RTs for the final successful pumps. For the latter measure, RTs below 100 ms and above 3000 ms were excluded following Kardos et al. (2016). No RT were excluded for the cash-out decisions. Bonferroni correction was used to account for multiple comparisons, resulting in a corrected significance threshold of $p < .025$ (.05/2). Since the assumption of normality was violated, Spearman's rank correlations were conducted. In addition, extreme outliers were removed prior to the analyses; participants were classified as extreme outliers if their values exceeded 3 × IQR below the first quartile (Q1) or above the third quartile (Q3).

First, we tested whether theta decrease was associated with RTs for the cash-out decisions following the final successful pumps. No significant correlation was observed ($r_s(39) = 0.282$, $p = .074$, Fig. S1A). Second, we examined whether theta decrease was related to RTs for the final successful pumps. A significant positive correlation was observed ($r_s(40) = 0.434$, $p = .004$, Fig. S1B). Because more negative theta values indicate stronger suppression, this association suggests that faster execution of the final successful pump was accompanied by stronger post-response theta suppression. This pattern is consistent with the interpretation that post-response theta suppression reflects reduced monitoring demands or increased confidence following decision commitment.

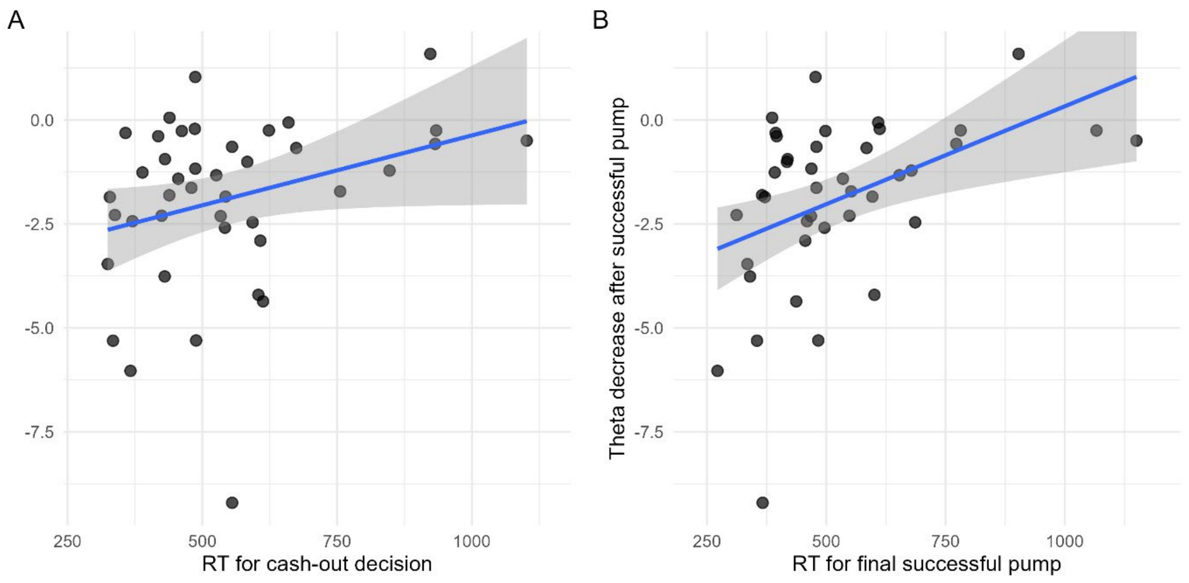


**Figure S1.** Associations between theta-band activity decrease following the final successful pumps and behavioral response times (RTs). The left panel shows the relationship between theta decrease and RTs for the cash-out decisions following the final successful pumps. The right panel shows the relationship between theta decrease and RTs for the final successful pumps. Each point represents an individual participant. Lines indicate linear regression fits with 95% confidence intervals. Theta decrease reflects mean total power in the theta band following the final successful pumps, with more negative values indicating stronger suppression. Correlation analyses were conducted using Spearman's rank correlation. No significant association was observed with the cash-out RTs, whereas a significant positive association was observed between theta decrease and RTs for the final successful pumps.
